# Profilin 1 Controls the Assembly, Organization, and Dynamics of Leading Edge Actin Structures Through Internetwork Competition and Collaboration

**DOI:** 10.1101/849356

**Authors:** Kristen Skruber, Peyton V. Warp, Rachael Shklyarov, James D. Thomas, Maurice S. Swanson, Jessica L. Henty-Ridilla, Tracy-Ann Read, Eric A. Vitriol

**Affiliations:** Department of Anatomy and Cell Biology, University of Florida, College of Medicine, Gainesville, FL, USA, 32610; Department of Molecular Genetics and Microbiology, Center for NeuroGenetics and the Genetics Institute, University of Florida, College of Medicine, Gainesville, FL, USA, 32610; Department of Cell and Developmental Biology, SUNY Upstate Medical University, NY, USA, 13210

## Abstract

How actin monomers are distributed to different networks remains poorly understood. One emerging concept is that the monomer pool is limited and heterogenous, causing biased assembly and internetwork competition. However, most knowledge regarding monomer distribution comes from studies where competing networks are discrete. In metazoans, many actin-based structures are complex, containing competing networks that overlap and are functionally interdependent. Addressing how monomers control the assembly and organization of these complex structures is critical to understanding how actin functions in cells. Here, we identify the monomer-binding protein profilin 1 (PFN1) as a major determinant of actin assembly, organization, and network homeostasis in mammalian cells. At the leading edge, PFN1 controls the localization and activity of the assembly factors Arp2/3 and Mena/VASP, with discrete stages of internetwork competition and collaboration occurring at different PFN1 concentrations. This causes substantial changes to leading edge actin architecture and the types of structures that form there.

## Introduction

To divide, move, and communicate, cells rely on a dynamic actin cytoskeleton that can rapidly assemble and adapt. This is achieved by the polymerization of actin monomers into filaments, the construction of large filament networks, and the disassembly of these networks back into monomers. To meet the demands of actin network assembly, cells maintain a large monomer reserve (1–4). However, several factors complicate how monomers are distributed to different actin structures within the cell (5). For example, monomers can undergo biased assembly into specific networks through interactions with polymerases and monomer-binding proteins (6, 7). Monomers can also be subcellularly localized (8, 9) or recycled back into the structures which they originated from (10). Actin isoforms can exhibit differential localization (11), dynamics (12), and regulation by post-translational modifications (13). Finally, the monomer/filament ratio is in homeostasis (14), which causes networks competing for the same monomers to alter their growth based on the activity or expression level of different assembly factors (6, 7, 14). Thus, the rules that govern how monomers are allocated throughout the cell are complicated and many of the details still need to be uncovered.

In cells, most actin monomers are bound to profilin (15). Profilin prevents spontaneous nucleation and directs monomers to the fast-growing (barbed) ends of actin filaments (15, 16). Profilin also interacts with formins (17–21) and Mena/VASP (7, 22, 23) at barbed ends and enhances their ability to polymerize actin. Although profilin has been shown to suppress branching of the multi-component assembly factor Arp2/3 (6, 7, 24, 25), it can supply monomers to Arp2/3-driven networks through interactions with WASP-family proteins (26–28). Thus formins, Mena/VASP, and Arp2/3 all utilize profilin-actin (19, 20, 22, 28–30), which can cause internetwork competition (6, 7). The concept of monomer competition has largely been inferred from experimental systems where actin networks are spatially and functionally distinct. These include in vivo experiments in yeast, where monomers polymerize into either Arp2/3-driven patches or formin-based cables (31). Metazoan cells, however, contain actin structures where complex filament architectures are constructed by multiple assembly factors. Two examples of this are the lamellipodia, a protrusive structure at the leading edge of cells which is made of dendritic and linear actin networks assembled by Arp2/3, formins, and Mena/VASP (32–35) and filopodia, the finger-like actin projections that extend beyond the lamellipodia (34). It is poorly understood how the monomer pool regulates the assembly and organization of these types of actin superstructures, where networks competing for monomers overlap and are functionally interdependent.

In this study, we dissect how actin assembly factors collectively construct complex actin networks through Profilin 1 (PFN1). Using a PFN1 knockout/rescue experimental paradigm, we demonstrate that PFN1 controls the majority of actin polymerization, determines the monomer to filament ratio set point, and facilitates homeostatic interplay between different actin networks. These properties extend to PFN1’s regulation of actin at the leading edge, where it coordinates the localization and activity of different actin assembly factors in a concentration-dependent manner and determines which leading edge structures assemble. Thus, monomer distribution through PFN1 is a major determinant of actin assembly, organization, and homeostasis in cells and a critical regulator of complex actin architectures.

## Results and Discussion

### PFN1 controls global actin polymerization and monomer/filament homeostasis

We knocked out the PFN1 gene with CRISPR/Cas9 in Cath.a differentiated (CAD) cells. CAD cells were chosen for this study because we have previously characterized how their monomer pool influences actin network behavior (8, 10, 12, 36) and because PFN1 is the predominant profilin isoform. PFN2 is expressed approximately 10-fold less than PFN1, while PFN3 and PFN4 are not expressed. Importantly, PFN2 expression doesn’t change upon deletion of PFN1 (Supplementary Table 1), which is consistent with studies showing that these isoforms cannot compensate for each other (37). Additionally, very few other actin-binding proteins are differentially expressed when PFN1 is knocked out (Supplementary Table 2). Thus, this is a strong cell line to study how PFN1 regulates the actin cytoskeleton. Loss of PFN1 was verified by western blot (Fig. 1A, B). The actin cytoskeleton is drastically altered in PFN1 KO cells, which can be rescued with physiologically relevant expression levels of GFP-PFN1 (Fig. 1A, B).

**Figure 1.**
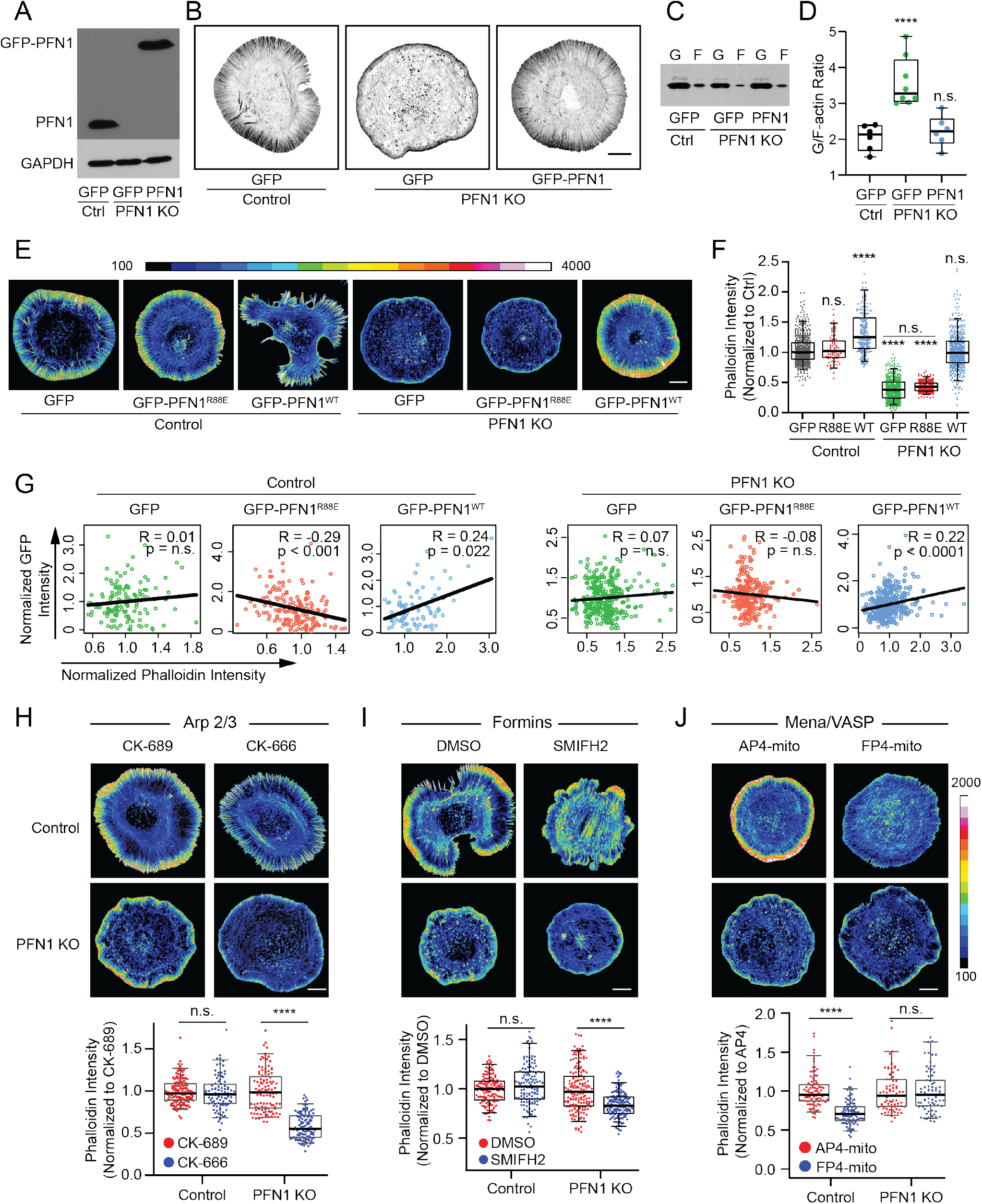
PFN1 controls global actin polymerization and monomer/filament homeostasis. **(A)** Western blot for profilin 1 (PFN1) of control (Ctrl) and PFN1 knockout (KO) CAD cells transfected with GFP or GFP-PFN1 (PFN1). **(B)** Representative images of the actin cytoskeleton in control and PFN1 KO cells transfected with GFP or GFP-PFN1. Actin filaments were labeled with Alexa568-phalloidin. Scale bar, 10 *μ*m. **(C)** Western blot for actin in control and PFN1 KO cells expressing GFP or GFP-PFN1. Cell lysates were subjected to ultracentrifugation to separate monomers and filaments. Actin monomers (G-actin, designated as ‘G’) remain in the supernatant while filaments (F-actin, designated as ‘F’) sediment to the pellet. (Legend continues on next page). **(D)** Quantification of the G/F-actin ratio from **(C)**. Individual data points are plotted along with the mean and 95% confidence intervals. Number of biological replicates is as follows: control + GFP (n = 6), PFN1 KO + GFP (n = 8), and PFN1 KO + GFP-PFN1 (n = 6). **(E)** Control and PFN1 KO cells transfected with GFP, GFP-PFN1 (GFP-PFN1^WT^), or the non-actin binding PFN1 mutant R88E (GFP-PFN1^R88E^). Actin filaments have been labeled with Alexa568-phalloidin. The images are scaled identically and pseudocolored based on the included lookup table to convey relative fluorescent intensities. Scale bar, 10 *μ*m. **(F)** Quantification of mean Alexa568-phalloidin intensity in control and KO cells expressing GFP, GFP-PFN1^R88E^ (R88E), or GFP-PFN1 (WT). Data are plotted relative to CTRL cells expressing GFP. For control cells expressing GFP, GFP-PFN^R88E^ or GFP-PFN1^WT^, n = 475, 166, and 206, respectively. For PFN1 KO cells expressing GFP, GFP-PFN^R88E^ or GFP-PFN1^WT^, n = 446, 206, and 443, respectively. **(G)** Correlation between GFP and phalloidin intensity for cells in **(E)**. Intensities were normalized to the mean of each data set. **(H-J)** Representative images and quantification of mean Alexa568-phalloidin intensity in control and PFN1 KO cells, where Arp2/3, formins, or Mena/VASP were inhibited. The images are scaled identically and pseudocolored based on the included lookup table to convey relative fluorescent intensities. Scale bar, 10 *μ*m. **(H)** Control and PFN1 KO cells were pretreated with control (CK-689) or Arp2/3 (CK-666) small molecule inhibitors for 1 hour prior to fixation and labeling with Alexa568-phalloidin. For control cells n = 154 for CK-689 and 93 for CK-666. For PFN1 KO cells n = 111 for CK-689 and 119 for CK-666. **(I)** Control and PFN1 KO cells were pre-treated with vehicle control (DMSO) or the pan-formin small molecule inhibitor SMIFH2 for 20 min prior to fixation and labeling with Alexa568-phalloidin. For control cells n = 140 for DMSO and n = 130 for SMIFH2. For PFN1 KO cells n = 160 for DMSO and n = 159 for SMIFH2. **(J)** Control and PFN1 KO cells were transfected with the mitochondria sequestering Mena/VASP construct FP4 mito, or its AP4 mito control for 16 hour prior to fixation and labeling with Alexa568-phalloidin. For control cells n = 89 for AP4 mito and n = 94 for FP4 mito. For PFN1 KO cells n = 79 for AP4 mito and n = 96 for FP4 mito. Box-and-whisker plots in D,F,H,I,J denote 95th (top whisker), 75th (top edge of box), 25th (bottom edge of box), and 10th (bottom whisker) percentiles and the median (bold line in box). p values plotted relative to control + GFP, unless otherwise indicated. **** indicates p ≤ 0.0001, *** p ≤ 0.001, n.s. = not significant (p > 0.05). p values were generated by ANOVA followed by Tukey’s post hoc test (comparison of ≥ 3 conditions).

We first sought to determine how actin polymerization and monomer/filament homeostasis are disrupted in the absence of PFN1. Western blot quantification of monomer and filament-containing cellular fractions revealed that the ratio of monomeric to polymerized actin is 3:1 in PFN1 KO cells (Fig. 1C, D), and can be rescued to the control cell ratio by expressing GFP-PFN1 (Fig. 1C, D). Quantitative image analysis of cells labeled with fluorescent phalloidin was then used to measure relative amounts of actin filaments. Filament levels in PFN1 KO cells were reduced more than 50% in comparison to control cells (Fig. 1E, F). This reduction was rescued by expression of wild-type GFP-PFN1 but not the non-actin binding R88E mutant (38) (Fig. 1E, F). While knocking out PFN1 caused a slight reduction in actin expression (Supplementary Fig. 1A), it was insufficient to explain the substantially larger decrease in polymerized actin levels in PFN1 KO cells (Fig. 1E, F) and the reciprocal increase in the monomer/filament ratio (Fig. 1C, D). Additionally, supplying PFN1 KO cells with more actin by expressing EGFP-β-actin did not increase actin polymerization (Supplementary Fig. 1B, C), verifying that filament levels are low because the monomers cannot polymerize in the absence of PFN1. The substantial inability of actin to polymerize in PFN1 KO cells is likely due to a combination of an increase in filament capping (39), inhibition of or decrease in barbed-end polymerase activity (20, 22), a decrease in the nucleotide exchange rate of actin (40), and an increase in sequestering by Thymosin β4 (41).

Overexpressing PFN1 in control cells caused an increase in actin filaments (Fig. 1E, F). We postulated that PFN1 concentration was the predominant factor which determined the monomer to filament ratio and that global actin polymerization would scale with PFN1 expression levels. To test this, we performed a correlation analysis between PFN1 expression and polymerized actin. In control and KO cells, there was a significant, positive, linear correlation between actin filament levels and PFN1 expression, but not with expression of GFP or GFP-PFN1^R88E^ (Fig. 1G). Interestingly, the R88E mutant had a slightly dominant negative effect on actin polymerization, most likely due to binding poly-L-proline residues but not actin (38). These results demonstrate that PFN1 is not only needed for actin to polymerize (Fig. 1C-F), but that its expression level determines the monomer/filament homeostasis set point.

Monomer/filament homeostasis has been shown in yeast, where inhibiting Arp2/3-driven actin patches causes a reciprocal increase of formin-based actin cables (6, 14). To determine if PFN1 was needed to maintain internetwork homeostasis, we measured polymerized actin levels after specific actin assembly factors were inhibited. As predicted by previous work (6, 7), global filament levels were maintained in control cells after inhibition of Arp2/3 or formins (Fig. 1H, I). However, inhibiting Arp2/3 or formins in PFN1 KO cells caused a significant reduction in actin filaments (Fig. 1H, I), revealing an inability to undergo compensatory network assembly. Furthermore, inducing loss of function of Mena/VASP proteins by targeting them to mitochondria (Supplementary Fig. 2) did not alter actin polymerization levels in PFN1 KO cells (Fig. 1J), but did reduce actin filaments in control cells. This result supports prior work demonstrating that Mena/VASP requires profilin-actin for its polymerase activity (22, 29). Because of the potential for compensatory mechanisms to occur in the time needed to express the FP4-mito construct, these experiments cannot be directly compared to the immediate inactivation of Arp2/3 and formin by small molecule inhibitors. However, these experiments do still reveal that PFN1 plays an integral role in the assembly of Arp2/3, formin, and Mena/VASP-based networks. Arp2/3 and formins can function independently of PFN1, but their internetwork monomer/filament homeostasis is not maintained unless PFN1 is present. Actin assembly by Mena/VASP is PFN1-dependent and not in homeostasis with Arp2/3 and formin-based networks, demonstrating an exclusive use of PFN1-actin that does not transfer to other the assembly factors.

### PFN1 controls assembly and organization of actin at the leading edge

Since PFN1 was shown to coordinate actin polymerization by Arp2/3, formins, and Mena/VASP, we wanted to determine how it controlled the assembly and organization of actin structures that depend on all three assembly factors: the lamellipodia (32–34) and filopodia (convergent elongation ref). PFN1 KO cells have a dramatically altered lamellipodia (Fig. 2A) with approximately three-fold less actin than control cells (Fig. 2B). Linescan analysis revealed that PFN1 KO lamellipodia are also smaller (Fig. 2C). Live cell imaging of cells expressing Lifeact-mRuby demonstrated that actin retrograde flow rates were reduced in PFN1 KO cells by approximately 50% (Fig. 2D, E). All lamellipodia phenotypes in PFN1 KO cells could be rescued by expressing GFP-PFN1^WT^ but not GFP-PFN1^R88E^ (Fig. 2A-E). One factor that drives retrograde flow rate is the number of actively polymerizing filaments (42). To confirm that actin polymerization was reduced at the leading edge of PFN1 KO lamellipodia, we labeled sites of active polymerization (43) (see Material and Methods for details) and found that PFN1 KO cells have approximately 50% fewer actively polymerizing filaments than control cells (Fig. 2F, G). Again, this is likely due to a decrease in barbed end polymerase activity and an increase in filament capping (39, 44).

**Figure 2.**
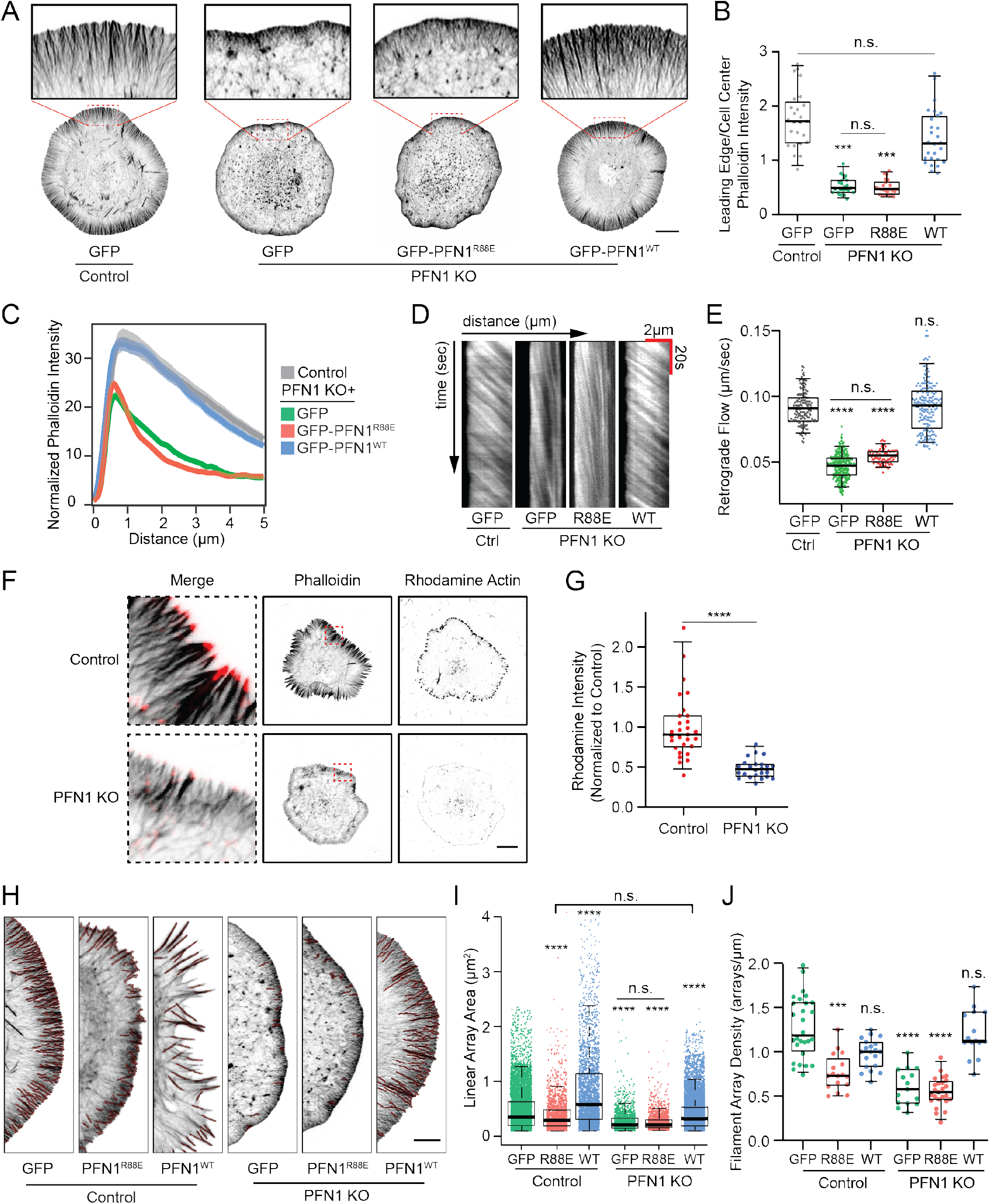
PFN1 controls assembly and organization of actin at the leading edge. **(A)** Representative confocal super-resolution images of control and PFN1 KO cells expressing GFP, GFP-PFN1^R88E^, or GFP-PFN1^WT^ and labeled with Alexa568-phalloidin. Insets highlight actin at the leading edge. Scale bar, 10 *μ*m. **(B)** Ratio between the sum intensity of the leading edge and the cellular interior of control and PFN1 KO cells expressing GFP, GFP-PFN1^R88E^ (R88E), or GFP-PFN1^WT^ (WT) and labeled with Alexa568-phalloidin. For all conditions n = 25. **(C)** Linescan analysis of the leading edge of control and PFN1 KO cells expressing GFP, GFP-PFN1^R88E^, or GFP-PFN1^WT^ labeled with Alexa568-phalloidin. The transparent bands depict 95% confidence intervals. For all conditions, n = 400 lines drawn from 20 cells. **(D)** Representative kymographs of lamellipodia retrograde flow in control and PFN1 KO cells expressing Lifeact-mRuby and either GFP, GFP-PFN1^R88E^ (R88E), or GFP-PFN1^WT^ (WT). **(E)** Quantification of retrograde flow from kymographs as depicted in **(D)**. For control cells n = 170 measurements from 17 cells. For PFN1 KO cells n = 310 measurements from 31 cells for GFP; n = 100 measurements from 10 cells for R88E; and n = 180 measurements from 18 cells for WT. **(F)** Confocal images showing actin filaments (Phalloidin) and barbed ends (Rhodamine Actin) in control and PFN1 KO cells. Cells were gently permeabilized and incubated with rhodamine actin in polymerization buffer for 60 s to label actin filament barbed ends. They were then fixed and incubated with Alexa488-phalloidin to label actin filaments. Black scale bar, 10 *μ*m. Red scale bar in inset, 1 *μ*m. **(G)** Quantification of barbed ends in control and PFN1 KO cells by measuring sum rhodamine actin fluorescence. Fluorescent intensity is plotted relative to control cells. For control cells n = 30, for PFN1 KO cells, n = 25. **(H–J)** Quantification of linear arrays at the leading edge of control and PFN1 KO cells expressing GFP, GFP-PFN1^R88E^ or GFP-PFN1^WT^. **(H)** Representative confocal super-resolution images of the leading edge of control and PFN1 KO cells expressing GFP, GFP-PFN1^R88E^, or GFP-PFN1^WT^ and labeled with Alexa568-phalloidin. Results from linear array segmentation analysis are outlined in red. Scale bar, 5 *μ*m. **(I and J)** Linear array area and density measurements of control cells expressing GFP (n = 5075 arrays, 28 cells), GFP-PFN1^R88E^ (n = 1937 arrays, 17 cells) or GFP-PFN1^WT^ (n = 2813 arrays, 20 cells) and PFN1 KO cells expressing GFP (n = 1559 arrays, 15 cells), GFP-PFN1^R88E^ (n = 1903 arrays, 25 cells) or GFP-PFN1^WT^ (n = 2688 arrays, 15 cells). Area measures the size of individual arrays. Density was measured by normalizing the linear array count to the cell perimeter. Box-and-whisker plots in B,E,G,I,J denote 95th (top whisker), 75th (top edge of box), 25th (bottom edge of box), and 10th (bottom whisker) percentiles and the median (bold line in box). p values plotted relative to control + GFP, unless otherwise indicated. **** indicates p ≤ 0.0001, *** p ≤ 0.001, n.s. = not significant (p > 0.05). p values were generated by either a two-tailed student’s t-test (comparison of two conditions) or by ANOVA followed by Tukey’s post hoc test (comparison of ≥ 3 conditions). For (I) Dunn’s post hoc test was used to generate p-values after normality of the data was assessed.

To understand how PFN1 expression alters lamellipodia architecture, we quantified linear filament arrays (see Materials and Methods for details), which are the predominant products of formins and Mena/VASP at the leading edge (33, 34, 45). In PFN1 KO cells, linear array area and density were both substantially reduced compared to control cells (Fig. 2H-J). The loss of linear arrays in PFN1 KO cells is rescuable by expressing GFP-PFN1^WT^ but not GFP-PFN1^R88E^ (Fig. 2H-J). PFN1 overexpression increased linear array area in control cells (Fig. 2H-J), demonstrating that the size of lamellipodia linear array networks is modulated by PFN1 concentration, in a manner similar to global actin polymerization (Fig. 1G). Interestingly, PFN1 overexpression caused the arrays to grow larger (Fig. 2I) but did not increase their density (Fig. 2J), suggesting that once a certain number of bundles has been assembled, these actin networks can only expand by increasing in size, perhaps due to space limitations and/or competition for resources from Arp2/3-based dendritic networks (14, 46).

### PFN1 coordinates Arp2/3 and Mena/VASP localization and activity at the leading edge

Next, we sought to understand how PFN1 controls Arp2/3 and Mena/VASP activity in the lamellipodia. Immunocytochemistry confirmed that Arp2/3 and Mena both localizedto the leading edge in control cells as expected (Fig. 3A-C). However, Arp2/3 is depleted from the leading edge of PFN1 KO cells and Mena localization is increased (Fig. 3A-C). We confirmed that this difference in leading edge localization cannot be explained by differential expression of Arp2/3 or Mena (Supplementary Fig. 3). This result was surprising as it has previously been shown that reducing PFN1 expression enhanced Arp2/3 localization to the leading edge (7), supporting a role for PFN1 as an inhibitor of Arp2/3 network assembly. However, in light of recent work showing that profilin transfers monomers to Arp2/3 networks via membrane-concentrated WASP-family proteins (28), we may be seeing a similar role for PFN1 in our experimental system that could only be revealed with a complete loss-of-function.

**Figure 3.**
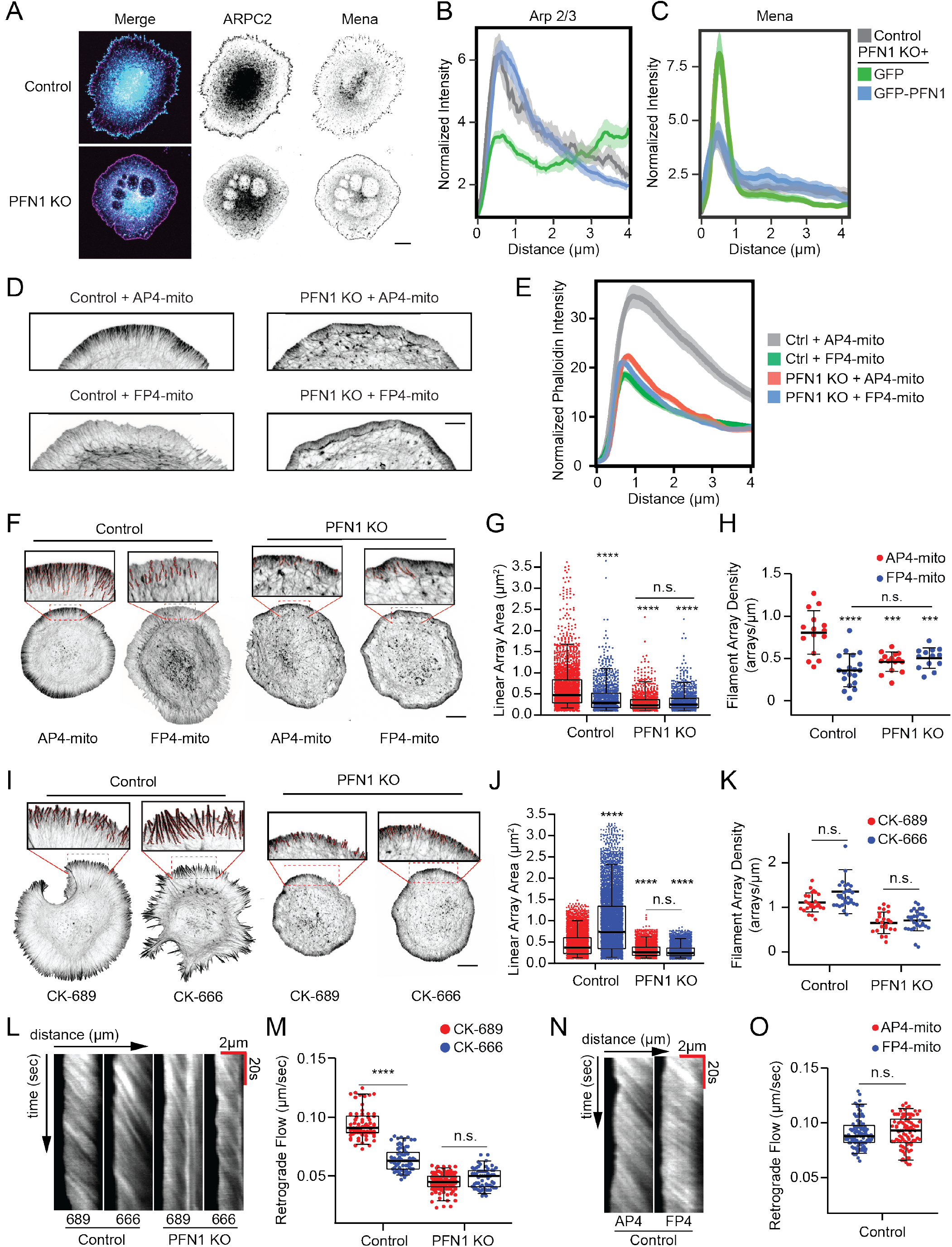
PFN1 coordinates Arp2/3 and Mena/VASP localization and activity at the leading edge. **(A)** Representative images of ARPC2 and Mena immunolabeling in control and PFN1 KO cells. Scale bar, 10 *μ*m. **(B)** Linescan analysis of ARPC2 immunolabeling at the leading edge of control and PFN1 KO cells expressing GFP or GFP-PFN1. The transparent bands depict 95% confidence intervals. For all conditions n = 300 linescans from 15 cells. **(C)** Linescan analysis of Mena immunolabeling at the leading edge of control and PFN1 KO cells expressing GFP or GFP-PFN1. The transparent bands depict 95% confidence intervals. For all conditions n = 400 linescans from 20 cells. **(D)** Representative confocal super-resolution images of the leading edge of control and PFN1 KO cells expressing AP4- or FP4-Mito and labeled with Alexa568-phalloidin. Scale bar, 5 *μ*m. **(E)** Linescan analysis of the leading edge of control and PFN1 KO cells expressing AP4- or FP4-Mito, and then labeled with Alexa568-phalloidin. For control cells expressing AP4-Mito, 320 lines were drawn from 16 cells; for control cells expressing FP4-Mito 400 lines were drawn from 20 cells; for PFN1 KO cells expressing AP4-Mito and FP4-Mito, 300 lines were drawn from 15 cells. The transparent bands depict 95% confidence intervals. **(F)** Representative confocal super-resolution images of the leading edge of control and PFN1 KO cells expressing the MENA/VASP mitochondria-sequestering FP4-mito or its control AP4-mito. Cells expressed the DNA vectors for 16 hrs before they were fixed and labeled with Alexa568-phalloidin. Results from linear array segmentation analysis are outlined in red. Scale bar, 10 *μ*m. **(G and H)** Linear array area and density measurements of control and PFN1 KO cells expressing AP4-mito or FP4-mito as depicted in **(F)**. For control + AP4-mito, n =1928 arrays, 15 cells; For control + FP4-mito, n = 962 arrays, 20 cells; For PFN1 + AP4-mito, n = 853 arrays, 15 cells; For PFN1 + AP4-mito, n = 961 arrays, 15 cells. **(I)** Representative confocal super-resolution images of the leading edge of control and PFN1 KO cells that were pre-treated with control (CK-689) or Arp2/3 (CK-666) small molecule inhibitors for 1 hr prior to fixation and labeling with Alexa568-phalloidin. Results from linear array segmentation analysis are outlined in red. Scale bar 10 *μ*m. **(J and K)** Linear array area and density measurements of control and PFN1 KO cells treated with CK-689 or CK-686 as depicted in **(I)**. For control + CK-689, n = 3846 arrays, 25 cells; For control + CK-666, n = 4777 arrays, 30 cells; for PFN1 KO + CK-689, n = 2078 arrays, 29 cells; for PFN1 KO + CK-666, n = 2856 arrays, 23 cells. Area measures the size of individual linear arrays. Density was measured by normalizing the linear array count to the cell perimeter. **(L)** Representative kymographs of lamellipodia retrograde flow in control and PFN1 KO cells expressing Lifeact-mRuby and treated with CK-689 or CK-666 for 60 minutes prior to imaging. **(M)** Quantification of retrograde flow from kymographs as depicted in **(L)**. For control + CK-689 and control + CK-666 n = 70 measurements, 7 cells; for PFN1 KO + CK-689 n = 80 measurements, 8 cells; for PFN1 KO + CK-666 n = 120 measurements, 12 cells. **(N)** Representative kymographs of lamellipodia retrograde flow in control cells expressing Lifeact-mRuby and either AP4-mito or FP4-mito. **(O)** Quantification of retrograde flow from kymographs as depicted in **(N)**. For control + AP4-mito n =100 measurements, 10 cells; for control + FP4-mito n = 90 measurements, 9 cells. Box-and-whisker plots in E,H, M,O denote 95th (top whisker), 75th (top edge of box), 25th (bottom edge of box), and 10th (bottom whisker) percentiles and the median (bold line in box). Data in F,I are plotted as median with interquartile range. p values plotted relative to control, unless otherwise indicated. **** indicates p ≤ 0.0001, *** p ≤ 0.001, n.s. = not significant (p > 0.05). p values were generated by either a two-tailed student’s t-test (comparison of two conditions) or by ANOVA followed by Tukey’s post hoc test (comparison of ≥ 3 conditions). For (G,J) Dunn’s post hoc test was used to generate p-values after normality of the data was assessed.

We have also shown that inducing loss of function by targeting Mena/VASP to mitochondria with FP4-mito (Supplementary Fig. 2) did not change total levels of Factin in PFN1 KO cells (Fig. 1G), suggesting that PFN1-actin is required for Mena/VASP polymerase activity. Thus, it was also perplexing that Mena concentrated at the leading edge in PFN1’s absence. Mena’s localization there could indicate Mena is tethered to the plasma membrane or localized to filament barbed ends (47). In the latter case, if Mena cannot function as a polymerase without profilin-actin (as indicated in Fig. 1G), it could be acting as a capping protein by preventing barbed end growth. To address these possibilities, we used linescan analysis of fluorescently labeled actin filaments to compare the intensity profiles in lamellipodia of control and PFN1 KO cells expressing FP4-mito. FP4-mito expression in control cells severely depleted lamellipodial actin in a manner that was nearly identical to knocking out PFN1, while it had no additional effect on the lamellipodia of PFN1 KO cells (Fig. 3D, E). Additionally, we measured linear arrays in the lamellipodia of FP4-mito expressing cells. As predicted, inhibiting Mena/VASP at the leading edge in control cells had significant effects on the size and density of linear arrays (Fig. 3F-H). However, there was no change in linear array properties when FP4-mito was expressed in PFN1 KO cells (Fig. 3F-H). Together, these data indicate that leading edge-localized Mena is inert in PFN1-depleted cells: it is not contributing to or inhibiting actin polymerization. These results also corroborate previous work showing that Mena requires profilin-actin for its polymerase activity (22, 29).

### Leading edge actin architecture and Arp2/3 activity are defined by PFN1 concentration

To determine if PFN1-mediated internetwork competition between Arp2/3, formins, and Mena/VASP drives complex actin network formation at the leading edge, we used electroporation to introduce defined amounts of PFN1 protein into cells. Protein electroporation can be performed on 10^6^ cells simultaneously with a >99% transfection rate. Additionally, by circumventing the normal biosynthetic pathway, proteins can be studied under conditions that minimize a transcriptional response (49, 50). Using fluorescently labeled dextran, we demonstrated that the amount of delivered protein was linearly proportional to the bath concentration of protein in the electroporation chamber. Moreover, the variability in electroporation efficiency was low, allowing for discrete concentrations of delivered material (Supplementary Fig. 4A-C). PFN1 protein could also be introduced into cells at defined concentrations, including physiological levels (Supplementary Fig. 4D,E). In the following section and in Figure 4, when we mention PFN1 concentration, we are referring to the bath concentration of the electroporation chamber and not the amount protein delivered into cells. Modulating the PFN1 concentration in PFN1 KO cells had dramatic effects on lamellipodia architecture. At 20 *μ*MPFN1, the lamellipodia was virtually eliminated and caused the cells to send out numerous filopodia protrusions (Fig. 4A, B). At intermediate concentrations (50 *μ*M) of PFN1, the lamellipodia returned and filopodia protrusions subsided (Fig. 4A, B). The 100 *μ*M concentration, which completely rescued PFN1 protein expression to control levels (Supplementary Fig. 4E), restored the size and architecture of the lamellipodia to strongly resembled control cells (Fig. 4A, B). Thus, the type and amount of actin filament structures that form at the leading edge are largely dependent on the availability of monomers bound to PFN1.

**Figure 4.**
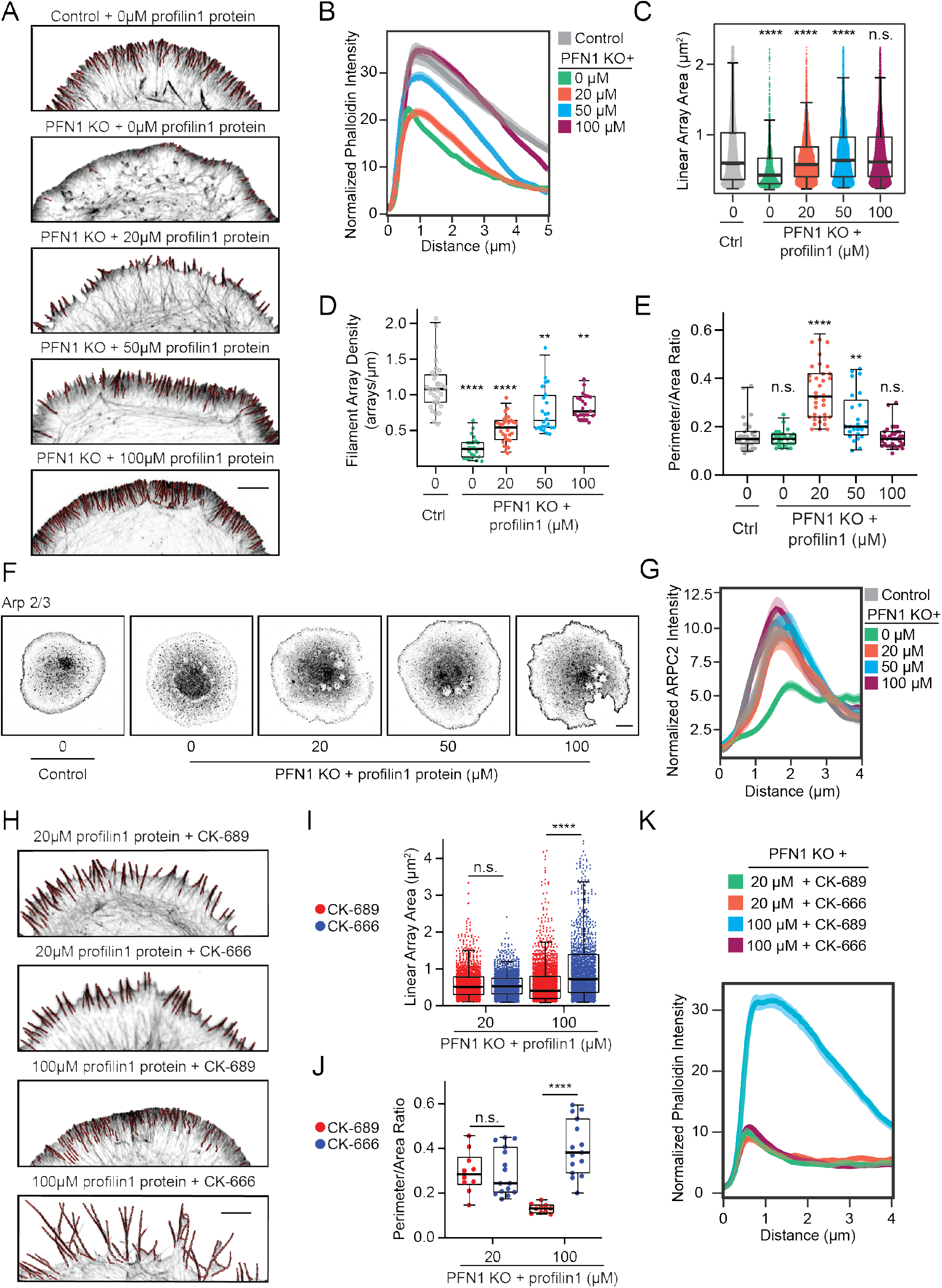
Lamellipodia architecture and Arp2/3 activity are defined by the cellular PFN1 concentration. **(A)** Representative confocal super-resolution images of the leading edge of control and PFN1 KO cells after electroporation with the designated concentration of purified PFN1 and then labeled with Alexa568-phalloidin. Insets highlight actin architecture at the leading edge. Scale bar, 5 *μ*m. **(B)** Linescan analysis of Alexa568-phalloidin fluorescence intensity at the leading edge of cells depicted in **(A)**. Only regions containing a lamellipodia were analyzed. The transparent bands depict 95% confidence intervals. For all conditions, n = 400 linescans from 20 cells. **(C and D)** Linear array area and density measurements of control and PFN1 KO cells electroporated with the designated concentration of purified profilin 1 as depicted in **(A)**. For control cells n = 5652 arrays, 27 cells. For PFN1 KO cells + 0 *μ*M n = 2609 arrays, 27 cells. For PFN1 KO cells + 20 *μ*M n = 3732 arrays, 34 cells. For PFN1 KO cells + 50 *μ*M n = 5265 arrays, 35 cells. For PFN1 KO cells + 100 *μ*M n = 4943 arrays, 30 cells. **(E)** Quantification of filopodia at the leading edge in control and PFN1 KO cells after electroporation with the designated concentration of purified PFN1 as depicted in **(A)**. Filopodia were quantified by taking a ratio of the cell perimeter to the cell area. Higher ratios reflect more filopodia protrusions. For control cells n = 35. For PFN1 KO cells electroporated with 0 *μ*M, 20 *μ*M, 50 *μ*M, and 100 *μ*M n = 29, 36, 25, and 34, respectively. **(F)** Representative confocal images of control and PFN1 KO cells electroporated with the indicated concentration of purified profilin 1 and immunolabeled for Arp2/3 with an ARPC2 antibody. Scale bar, 10 *μ*m. **(G)** Linescan analysis of ARPC2 fluorescence intensity at the leading edge of cells depicted in **(F).** The transparent bands depict 95% confidence intervals. For all conditions, n = 400 linescans from 20 cells. **(H)** Representative confocal super-resolution images of the leading edge of PFN1 KO cells after they were electroporated with the designated concentration of purified profilin 1, treated with CK-689 or CK-666 for 60 minutes, and then labeled with Alexa568-phalloidin. Scale bar, 5 *μ*m. **(I)** Linear array area measurements of PFN1 KO cells electroporated with 20 or 100 *μ*M of PFN1 protein and then treated with CK-689 or CK-686 as depicted in **(H)**. For PFN1 KO + 20 *μ*M PFN1, CK-689, n =1329 arrays, 15 cells; For PFN1 KO + 20 *μ*M PFN1 + CK-666, n = 1511 arrays, 10 cells; for PFN1 KO + 100 *μ*M profilin 1, CK-689 n = 1790 arrays, 10 cells; for PFN1 KO + 100 *μ*M PFN1, CK-666, n = 1424 arrays, 10 cells. Area measures the size of each individual linear array. **(J)** Quantification of filopodia at the leading edge in PFN1 KO cells after electroporation with 20 or 100 *μ*M of profilin 1 protein and then treatment with CK-689 or CK-686 as depicted in **(H)**. Filopodia were quantified by taking a ratio of the cell perimeter to the cell area. Higher ratios reflect more filopodia protrusions. For PFN1 KO cells electroporated with 20 *μ*M n=10, 15 for cells treated with CK-689 and CK-66, respectively. For PFN1 KO cells electroporated with 100 *μ*M n = 10, 15 for cells treated with CK-689 and CK-666, respectively. **(K)** Linescan analysis of phalloidin fluorescence intensity at the leading edge of cells depicted in **(H).** The transparent bands depict 95% confidence intervals. For all conditions, n = 200 linescans from 10 cells. Lines were drawn in between arrays. Box-and-whisker plots in C,D,E,J,K denote 95th (top whisker), 75th (top edge of box), 25th (bottom edge of box), and 10th (bottom whisker) percentiles and the median (bold line in box). p values plotted relative to control + 0 *μ*M profilin 1 protein, unless otherwise indicated. **** indicates p ≤ 0.0001, *** p ≤ 0.001, n.s. = not significant (p > 0.05). p values were generated by ANOVA followed by Tukey’s post hoc test (comparison of ≥ 3 conditions). For (C,I) Dunn’s post hoc test was used to generate p-values after normality of the data was assessed.

Linear array analysis of the leading edge of PFN1 KO cells rescued with varying amounts of PFN1 further revealed how PFN1 controls leading edge actin architecture. Interestingly, the size and density of linear arrays increases with PFN1 concentration (Fig. 4E-G). However, at lower PFN1 concentrations, the linear arrays largely exist as filipodia-like protrusions. At higher PFN1 concentrations, these protrusions subside and the linear arrays exist primarily as filament bundles within the lamellipodia network (Fig. 4E). Measuring the cell’s perimeter/area ratio, which is significantly increased when filopodia are present, confirmed the biphasic regulation of filopodia by PFN1 concentration (Fig. 4H). Along with previous experiments (Fig. 4A-D), this highlights the sensitivity of leading edge actin architecture to the availability of PFN1-actin. Since Arp2/3 localization at the leading edge is severely reduced in PFN1 KO cells (Fig. 3A, B) and intermediate concentrations of PFN1 favor formation of filopodia over lamellipodia (Fig. 4A, B), we wanted to determine how PFN1 concentration affects Arp2/3 localization to the leading edge. Surprisingly, we found that the majority of Arp2/3 returns to the leading edge upon introduction of a low (20 *μ*M) PFN1 concentration (Fig. 4C, D), despite no change in lamellipodia size in comparison to PFN1 KO cells (Fig. 4B). While Arp2/3 localization continued to increase with higher PFN1 concentrations (Fig. 4C, D), 20 *μ*M was sufficient to recall the majority of Arp2/3 back to the leading edge.

Discovering a PFN1 concentration that potently stimulated the production of filopodia-like protrusions allowed us to test whether their formation was dependent on Arp2/3 activity, as suggested by the convergent elongation model (51). This model proposes an interdependence of filopodia generation on Arp2/3-dependent nucleation, where Arp2/3 generates the base of the filament (51, 52). To test for convergent elongation, we combined controlled PFN1 delivery with CK-666-mediated inhibition of Arp2/3 (Fig. 4H-K). When PFN1 concentration was limited to cause a strong induction of filopodia, there was no difference in leading edge architecture after Arp2/3 inhibition, demonstrating these filipodia are Arp2/3-independent. Through Arp2/3 inhibition, we also show that the filopodia seen at low PFN1 concentrations (Fig. 4A,H) are not the result of the dissolution of a dendritic network and subsequent unmasking of stable filopodia (53), but rather due to the selective polymerization of specific structures. At 20 □M PFN1, CK-666 had no effect on linear filament arrays at the leading edge (Fig. 4 I,J). Arp2/3 inhibition similarly had no effect on lamellipodia actin as measured by linescan analysis of fluorescently-labeled actin filaments (Fig. 4H,I). However, when cells were given 100 *μ*M PFN1 and treated with CK-666, the actin structures that form are highly affected by Arp2/3 inactivation: the lamellipodia is severely decreased (Fig. 4K) and large filopodia form (Fig. 4I, J), demonstrating a shift toward assembly through formins and Mena/VASP. These data corroborate other studies showing PFN1 is crucial for homeostasis upon Arp2/3 inhibition (6, 7) but demonstrate an additional level of regulation where Arp2/3 activity also relies on PFN1.

PFN1 KO cells are affected by Arp2/3 inactivation (Fig. 1H) indicating that Arp2/3 globally contributes to building actin networks without PFN1. Although at the leading edge, Arp2/3 localization (Fig. 4F,G) and activity (Fig. 3L,M) require PFN1. Polymerization of linear filamentarrays by PFN1, formins, and Mena/VASP (Fig. 3F-K) may be necessary to create Arp2/3 binding sites at the leading edge, defining a collaboration between the different networks. This is reminiscent of filipodia/veil motility first identified in neuronal growth cones, where filopodia provide the initial step of membrane protrusions, followed by Arp2/3-based dendritic networks (54). However, low concentrations of PFN1 favor linear array network assembly (Fig. 4A-E), and prevent Arp2/3 from assembling dendritic networks (Fig. 4H-K). Dendritic networks may rely less heavily on PFN1 than linear networks do and thus may not be able to compete for profilin-actin. Higher PFN1 concentrations reduce this competition and allow all networks to form, though the homeostatic setpoint can be modulated by increasing PFN1 (Fig. 3I-K, 4H-K, 100 *μ*m). This result helps reconcile studies where PFN1 has been shown to both inhibit (7) and enhance (28) Arp2/3-based network assembly. Thus, by carefully controlling protein concentration, we were able to determine how PFN1 distributes monomers to different networks, even within complex actin structures. We speculate that PFN1 expression levels will largely determine the types of actin assembly that occur in a cell. Further, we have demonstrated that the types of actin structures that form within a cell can be dictated solely by PFN1 availability. Thus, a cell could dramatically change its actin networks by modulating PFN1 gene expression or by using phosphorylation to reduce its ability to bind actin (55).

## Methods

### DNA constructs

The following DNA constructs were used in this study: EGFP-PFN1 (Plasmid #56438, Addgene), Lifeact mRuby (pN1-Lifeact-mRuby, provided by Roland Wedlich-Soldner, Max-Planck Institute of Biochemistry), pEGFP-C1 EGFP β-actin, pMSCV EGFP-FP4 mito and EGFP-AP4 mito (provided by Alpha Yap) (33, 47). EGFP-PFN1R88E was generated from EGFP-PFN1 with site-directed mutagenesis (Q5 New England Biolabs) using the primers AATGGATCTTGAAACCAAGAGCACC (forward) and GTAAATTCCCCG-TCTTGC (reverse). Mutagenesis was confirmed by sequencing (Genewiz). EGFP-ß-actin was generated in a previous study (10). PFN1 KO cells were generated with the pCRISPR-CG02 vector (Genecopoeia) containing an sgRNA targeting TCGACAGCCTTATGGCGGAC in the mouse PFN1 gene and the puromycin donor plasmid pDonor-D01 (Genecopoeia) for selection. Control knock-out cells were generated using the same vectors and a scrambled sgRNA control targeting the sequence GGCTTCGCGCCGTAGTCTTA. All constructs were prepared for transfection using the GenElute HP Endotoxin-Free Plasmid Maxiprep Kit (Sigma-Aldrich).

### RNA-seq analysis

Total RNA was extracted from wild-type PFN1 knockout cells (four biological replicates per condition) and total RNA was used as input to generate strand-specific, rRNA-depleted RNA-seq libraries using the KAPA stranded RNA-seq Kit with RiboErase HMR (Kapa Biosystems). All steps were performed according to the manufacturer’s protocol except for the use of custom Illumina-compatible index primers to allow multiplexing. Paired-end, 36 bp sequencing of the final libraries was performed using an Illumina NextSeq500. Gene expression analysis was performed as previously described (56). Briefly, reads were de-multiplex based on sample specific barcodes and mapped to the human genome (hg19) using OLego (57). Uniquely mapped reads were assigned to genomic features and counted using Quantas (58). TMM normalization and identification of differentially expressed genes ws computed using edgeR (59). Final gene lists were filtered (|log2 fold change| ≥ 1; adjusted P ≤ 0.01) to identify significant changes.

### Cell culture

Cath.-a-differentiated (CAD) cells (purchased from Sigma-Aldrich) were cultured in DMEM/F12 medium (Gibco) supplemented with 8% fetal calf serum, 1% L-Glutamine, and 1% penicillin-streptomycin. Prior to imaging, CAD cells were plated on coverslips coated with 10 *μ*g/mL Laminin (Sigma-Aldrich). DMEM/F12 medium without phenol red (Gibco) supplemented with 15mM HEPES was used for live-cell imaging. CAD cells are a unique mouse neuroblastoma cell line that differentiate into a neuronal-like cell morphology upon serum withdrawal (60). We routinely use serum withdrawal to validate CAD cells by ensuring that they are able to undergo neuronal differentiation as evidenced by the formation of long (> 100 *μ*m), narrow projections after 2 days. Cell lines were also routinely tested for mycoplasma using the Universal Detection Kit (ATCC).

PFN1 KO cells were generated with CRISPR/Cas9 by transfecting CAD cells with the constructs described above. One week after transfection, cells that were modified by CRISPR/Cas9 were selected with 10 *μ*g/mL puromycin (Santa Cruz Biotechnology). This concentration was chosen as it kills 100% of cells that do not have the puromycin resistance gene within 24 hours (10, 12, 36). Puromycin was removed 24 hours prior to experiments that required transfection.

### Protein purification

Human profilin 1 protein was purified as described (61). Briefly, profilin 1 plasmids were cloned between NdeI and EcoRI sites of pMW172, a pET derivative (62, 63) and were expressed in Rosetta pRARE2 BL21(DE3) cells. Cells were grown in terrific broth to OD600 = 0.5 at 37 °C, then induced with IPTG for 3 h at 37 °C. Pellets were resuspended in 50 mM Tris HCl (pH 8.0), 10 mg/mL DNase I, 20 mg/mL PMSF, 1× protease inhibitor cocktail, and 10 mM DTT. Cells were lysed with 150 mg/mL lysozyme and sonicated at 4 °C. The cell lysate was clarified by centrifugation at 20,000 × g. The supernatant was passed over at QHighTrap column (GE Healthcare, Marlborough, MA) equilibrated in 50 mM Tris-HCl (pH 8.0), 1 M KCl, 10 mM DTT and the flow-through (containing PFN1) was collected and then applied to a Superdex 75 (10/300) gel filtration column (GE Healthcare) equilibrated in 50 mM Tris (pH 8.0), 50 mM KCl, 10 mM DTT. Fractions containing Profilin were pooled, aliquoted, and stored at 80 °C. Thawed Profilin aliquots were pre-cleared at 279,000 × g before use.

### DNA and protein electroporation

The Neon Transfection System (Invitrogen) was used to introduce DNA constructs and purified protein into cells using the 10 *μ*L transfection kit. Briefly, cells were allowed to reach a confluency of 70-80%, trypsinized and pelleted by centrifugation. The pellet was rinsed with DPBS and resuspended in a minimum amount of buffer R (Invitrogen) with a total of 1μg of DNA or the designated concentration of protein. Cells transfected with DNA constructs were given 14-18 hours after transfection before further experimental procedures were performed. For the 0 *μ*M concentration in protein transfections, cells were transfected with an equivalent amount of protein buffer. For protein electroporation, cells were given 2.5 hours to adhere on laminin coated coverslips before experiments were performed. In the combined protein electroporation and Arp2/3 inhibition experiment, cells were given 1 hour to adhere, media was changed and cells were treated for an additional hour with CK-666. A single 1400 v 20 ms pulse was used for both DNA and protein electroporation. This protocol routinely gave > 99% transfection efficiency.

### Western blots

Adherent cells were harvested with a cell scraper in RIPA buffer with cOmplete EDTA-free Protease Inhibitor Cocktail Roche (Millipore Sigma). Whole cell lysates were prepared by membrane disruption using repeated passage through a 27 gauge needle. Protein content was then assessed with Pierce BCA Protein Assay Kit (Thermo Fisher) and diluted in SDS buffer stained with Orange G (40% glycerol, 6% SDS, 300 mM Tris HCl, pH 6.8). 10μg samples were evenly loaded on an SDS-PAGE gel (Novex 4-20% Tris-Glycine Mini Gels, Thermo Fisher, or 15% gel as indicated). Protein was transferred to a PVDF membrane (0.2 micron, Immobilon) and blocked in 5% Bovine Serum Albumin (BSA) (Sigma-Aldrich) for 20 mins. All antibodies were diluted in 5% BSA and 0.1% Tween-20 (Fisher Scientific). Primary antibodies were incubated at 4°C overnight and secondary antibodies (Li-Cor; Abcam) were incubated for 2 hours at room temperature. Actin, profilin and GAPDH from whole cell lysate were detected with Li-Cor fluorescent antibodies on an Odyssey detection system (Li-Cor) or via X-ray film after incubation with a developing reagent (Thermo Fisher), as indicated. WesternSure Pre-Stained Chemiluminescent Protein Ladder (Li-Cor) was used as a molecular weight marker. The following antibodies/dilutions were used: rabbit anti-Profilin-1 (C56B8, 1:1000 dilution, Cell Signaling Technology); rabbit anti-Pan Actin (4968, 1:1000 dilution, Cell Signaling Technology); rabbit anti-GAPDH (2118, 1:3000 dilution, Cell Signaling Technology), goat anti-rabbit Alexa Fluor™ 680 (Li-Cor) was used at 1:3500 dilution for imaging on the Li-Cor Odyssey detection system and goat anti-rabbit HRP (Abcam) was used for X-ray detection.

### Actin monomer/filament ratio measurements

Cells were collected in lysis and F-actin stabilization buffer (LAS01, Cytoskeleton Inc) at 37°C in the presence of Halt Protease Inhibitor Cocktail (Thermo Fisher) and 10mM ATP (Cytoskeleton Inc). Cells were harvested via cell scraper and incubated at 37°C for 10 minutes. Unbroken cells and debris were pelleted at room temperature at 250 g for 3 mins and the supernatant was then immediately centrifuged at 150,000 g at 37°C for 1 hour in a swinging bucket rotor. The supernatant was carefully removed and the pellet was resuspended in a volume of F-actin depolymerization buffer (FAD-02, Cytoskeleton Inc.) matching the volume of the supernatant. All samples were then incubated on ice for 1 hour with periodic trituration and SDS buffer stained with Orange G (40% glycerol, 6% SDS, 300 mM Tris HCl, pH 6.8) was then added to each sample. Samples were then analyzed by western blot.

### Microscopy

Most images were acquired with a Nikon A1R+ laser scanning confocal microscope with a GaAsP multidetector unit. The microscope is also equipped with total internal reflection fluorescence (TIRF) and an ORCAFlash 4.0 sCMOS camera. All confocal and TIRF imaging was performed with an Apo TIRF 60X 1.49 NA objective. Deconvolution-based super-resolution confocal microscopy(64) (Wilson, 2011) was performed by using zoom settings higher than the Nyquist criteria, resulting in oversampled pixels (0.03 *μ*m). Confocal z-stacks were created and then deconvolved with Nikon Elements software using the Landweber algorithm (15 iterations, with spherical aberration correction) to create images with approximately 150 nm resolution (65). Live cell imaging was performed using TIRF. All cells analyzed for retrograde flow were non-motile. For live cell imaging, a stage incubator with CO2 and temperature control (Tokai Hit) was also used. Low resolution images used for measuring total actin levels (Fig. 1) and electroporation efficiency (Fig. S4) were taken with an EVOS XL digital inverted microscope objective (Life Technologies) equipped with a Plan Neoflour 20X 0.5 N.A. objective.

### Pharmacological inhibition of actin binding proteins

To inhibit Arp2/3, cells were treated with 50 *μ*M of the Arp2/3 inhibitor CK-666 or its control analog CK-689 (Sigma-Aldrich) for 1 hour prior. Stock solutions of 40 mM CK-666 and 100mM CK-689 were prepared in DMSO. To inhibit formins, cells were then treated with 10μM SMIFH2 (Sigma-Aldrich) or DMSO for 30 minutes. A stock solution of 25 mM SMIFH2 was prepared in DMSO.

### Immunofluorescence

Cells were fixed with 4% electron microscopy grade paraformaldehyde (Electron Microscopy Sciences) for 10 min at RT and then permeabilized for 3 minutes with 0.1% Tween-20. Cells were then washed three times with PBS and stained overnight at 4°C with primary antibodies diluted in PBS. They were then washed twice with PBS for 5 min, incubated with secondary antibodies (diluted 1:1000) for 1 hr at room temperature in PBS. Actin filaments were stained with Alexa Fluor 488 phalloidin or Alexa Fluor 568 phalloidin (diluted 1:100, Life Technologies) for 30 min at room temperature in immunofluorescence staining buffer. Cells were washed three times with PBS before mounting with Prolong Diamond (Life Technologies). The following antibodies were used: Rabbit anti-ARPC2 (p34-Arc, EMD Millipore) and mouse anti-Mena (clone A351F7D9, EMD Millipore) were used at a 1:500 dilution, anti-mouse IgG 647 and anti-rabbit IgG 568 (Life Technologies) were used at 1:1000 dilution.

### Fluorescently labeling sites of active polymerization

To label sites of active polymerization, cells were plated on coverslips that were pre-incubated with 10 *μ*g/mL Poly-D-Lysine (Sigma-Aldrich) for one hour and then washed with PBS prior to laminin coating. Barbed ends were labeled using a protocol adapted from (66). Briefly, a stock solution of permeabilization buffer (20 mM HEPES, 138 mM KCl, 4 mM MgCl2, 3 mM EGTA, and 1% BSA, pH 7.4) was prepared. Immediately prior to use, 0.025% saponin and 1 mM ATP were added followed by 0.45 *μ*M rhodamine actin (Cytoskeleton Inc). The culture medium was carefully removed by pipette and enough permeabilization buffer was added to cover cells. After 1 minute, permeabilization buffer was gently removed by pipette, rinsed briefly with 1X PBS and immediately fixed in 4% paraformaldehyde in cytoskeleton stabilization buffer (67) for 10 min. After carefully washing in 1X PBS 3 times, samples were incubated with phalloidin-488 or phalloidin-568 (diluted 1:100, Life Technologies) for 20 min at room temperature. Cells were imaged using identical conditions for comparison.

### Image Analysis

#### Quantification of total actin filaments per cell

Cells were transfected with GFP or a GFP-PFN1 construct, and then fixed and labeled with Alexa568-phalloidin. Images were taken on the EVOS XL microscope using identical illumination and camera exposure conditions. Image files were imported into ImageJ, the background was subtracted, and cells were segmented by fluorescence intensity-based thresholding using the phalloidin channel. Mean GFP and Alexa568-phalloidin values were used to assess relative PFN1 and F-actin levels in each cell. At least three biological replicates were performed per condition.

#### Quantification of retrograde flow

Cells transfected with Lifeact-mRuby were plated onto laminin-coated coverslips for 90 min. and then imaged with TIRF microscopy at 1 frame/s. Images were exported into ImageJ for analysis. Ten kymographs were generated per cell at the leading edge where fiduciary markers were clearly visible. Retrograde flow was calculated by measuring the distance and time that fiduciary markers traveled in the kymograph and solving for rate. Two-three biological replicates were performed per condition.

#### Quantification of actin and actin-binding proteins in the lamellipodia

Cells labeled with Alexa568-phalloidin or immunolabeled for Arp2/3 and Mena were imaged using confocal microscopy. Images were exported into ImageJ for analysis. A maximum intensity projection was made for each confocal z-stack. Lines ten pixels in width were drawn perpendicular to the cell edge, and fluorescence intensity was measured along the line. 20 lines were drawn per cell. Three biological replicates were performed per condition.

#### Quantification of linear arrays in the lamellipodia

Cells labeled with Alexa568-phalloidin were imaged using confocal deconvolution super-resolution microscopy. Images were deconvolved using Nikon Elements software and then imported into ImageJ for analysis. Confocal z-stacks were converted into a single maximum intensity projection image and the lamellipodia of the cell was manually thresholded. The “Tubeness” ImageJ plugin (68) was used for linear array segmentation on the thresholded lamellipodia. Images were convolved with a sigma value three times the minimum voxel separation. The convolved image was binarized and the Analyze Particle function was used to threshold objects with low circularity (0.0-0.3) and an area measurement greater than 0.1 *μ*m. Filament density was calculated by normalizing the number of segmented arrays to the perimeter of the cell. Three biological replicates were performed per condition.

### Statistical Analysis

Statistical significance was assessed by either a twotailed student’s t-test (comparison of two conditions) or by ANOVA followed by Tukey’s post hoc test (comparison of three or more conditions) using Graphpad Prism software. Dunn’s post hoc test was used in place of Tukey’s with nonparametric datasets, as indicated.

## Acknowledgements

We would like to thank Dr. Alpha Yap and Suzie Verma for the generous gift of the EGFP-FP4-Mito and EGFP-AP4-Mito DNA constructs. This project was supported by a National Institutes of Health (NIH) Maximizing Investigators’ Research Award for Early Stage Investigators (R35GM133485) to J.L.H.-R. and an NIH Pathway to Independence Award (R00 NS087104) to E.A.V.

