## Supplementary Materials for "Profilin 1 Controls the Assembly, Organization, and Dynamics of Leading Edge Actin Structures Through Internetwork Competition and Collaboration"

| Gene | Symbol | logFC | PValue |
|---|---|---|---|
| Arp2/3 Complex subunit 1b | Arpc1b | 0.290 | 1.6E-07 |
| Arp2/3 Complex subunit 2 | Arpc2 | 0.056 | 2.7E-01 |
| Arp2/3 Complex subunit 3 | Arpc3 | 0.103 | 9.5E-02 |
| Arp2/3 Complex subunit 4 | Arpc4 | 0.102 | 4.3E-02 |
| Arp2/3 Complex subunit 5 | Arpc5 | 0.167 | 1.0E-03 |
| Capping Actin Protein | Capza1 | -0.109 | 2.3E-01 |
| Capping Actin Protein | Capza2 | -0.129 | 3.5E-02 |
| Capping Actin Protein | Capzb | -0.036 | 4.5E-01 |
| Cell Division Cycle 42 | Cdc42 | 0.038 | 4.5E-01 |
| Cofilin-1 | Cfl1 | -0.212 | 1.2E-05 |
| Calponin-1 | Cnn1 | 0.272 | 8.6E-01 |
| Coronin 1A | Coro1a | 0.126 | 3.8E-01 |
| Coronin 1B | Coro1b | 0.115 | 2.6E-02 |
| Cortactin | Cttn | 0.147 | 4.4E-03 |
| Dishevelled Associated Activator of Morphogenesis 1 | Daam1 | 0.196 | 7.7E-03 |
| Diaphanous Related Formin 1 | Diap1 | 0.379 | 5.1E-08 |
| Diaphanous Related Formin 2 | Diap2 | 0.255 | 7.7E-03 |
| Diaphanous Related Formin 3 | Diap3 | 0.028 | 6.1E-01 |
| Destrin, actin depolymerizing factor | Dstn | 0.187 | 1.7E-04 |
| Enah | Enah | -0.051 | 3.4E-01 |
| Enah/Vasp-Like | Evl | -0.165 | 3.8E-03 |
| Ezrin | Ezr | -0.062 | 3.6E-01 |
| Formin Homology 2 Domain Containing 1 | Fhod1 | 0.431 | 1.9E-06 |
| Formin Like 1 | Fmnl1 | -0.033 | 5.9E-01 |
| Formin Like 2 | Fmnl2 | 0.061 | 2.7E-01 |
| Formin Like 3 | Fmnl3 | -0.077 | 2.6E-01 |
| Gelsolin | Gsn | 0.777 | 3.7E-17 |
| Inverted formin-2 | Inf2 | 0.456 | 8.7E-12 |
| Junction Mediating And Regulatory Protein | Jmy | 0.063 | 3.4E-01 |
| Myocardin Related (MAL) | Mkl1 | 0.088 | 0.15679 |
| Profilin-2 | Pfn2 | 0.228 | 2.9E-05 |
| Tropomyosin1 | Tpm1 | -0.120 | 8.2E-02 |
| Twinfilin-1 | Twf1 | -0.137 | 3.4E-02 |
| Twinfilin-2 | Twf2 | 0.154 | 5.8E-02 |
| Vasodilator Stimulated Phosphoprotein | Vasp | -0.004 | 9.4E-01 |
| Vinculin | Vcl | -0.261 | 7.2E-07 |
| Vimentin | Vim | 0.238 | 1.4E-06 |
| Wiskott-Aldrich syndrome protein family member 1 | Wasf1 | -0.240 | 6.9E-05 |
| WD Repeat Domain 1 | Wdr11 | 0.026 | 0.66484 |
| Zyxin | Zyx | 0.184 | 9.8E-03 |

**Table S1. Genes of actin binding proteins with no expression change between control and PFN1 KO CAD cells, related to Figure 1.** Criteria for differential expression of genes: logFC >±0.5 and a p value of <0.0001

| Gene | Symbol | logFC | PValue |
|---|---|---|---|
| Cordon-bleu WH2 repeat | Cobll1 | +0.89 | 1.45E-47 |
| Gelsolin | Gsn | +0.78 | 3.67E-17 |
| Was/Wasl 1 | Wipf1 | +0.71 | 1.07E-12 |
| Arp2/3 Complex subunit 5-like | Arpc5l | +0.56 | 1.05E-18 |
| Villin-1 | Vil1 | +0.55 | 4.84E-11 |
| Formin Homology 2 Domain Containing 3 | Fhod3 | +0.55 | 1.11E-04 |
| β-actin | Actb | -0.49 | 3.29E-26 |
| Dishevelled-associated activator of Morphogenesis | Daam2 | -0.53 | 3.38E-23 |
| Thymosin β-4 | Tmsb4x | -0.94 | 7.63E-49 |
| Gamma-actin | Actg1 | -1.21 | 1.50E-39 |

**Table S2. Differentially expressed genes of actin binding proteins between control and PFN1 KO cells, related to Figure 1.** Criteria for differential expression of genes: logFC >±0.5 and a p value of <0.0001

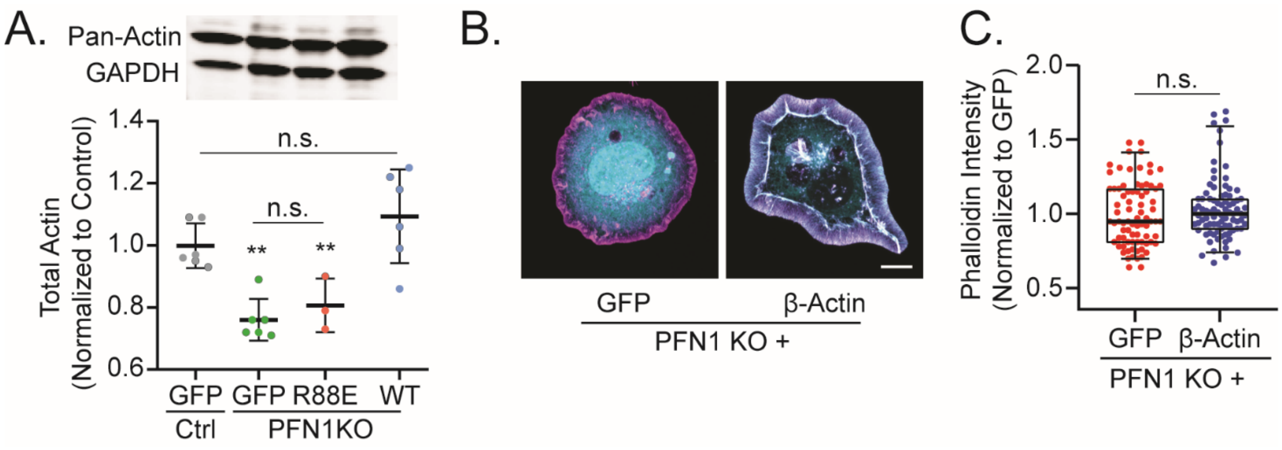


**Figure S1. Disruption of actin monomer/filament ratio in PFN1 KO cells is not due to decreased actin gene expression, related to Figure 1. (A)** Western blot analysis and quantification of β/γ-actin expression in control and PFN1 KO cells normalized to GAPDH as a loading control. n = 6 for control + GFP, PFN1 KO + GFP, $PFN^{WT}$ and n = 3 for PFN1 KO + $PFN^{R88E}$. **(B)** Effect of GFP-β-Actin expression on phalloidin intensity in PFN1 KO cells. n = 80 for GFP, 86 for GFP-β-Actin. Scale bar is 10 μm.

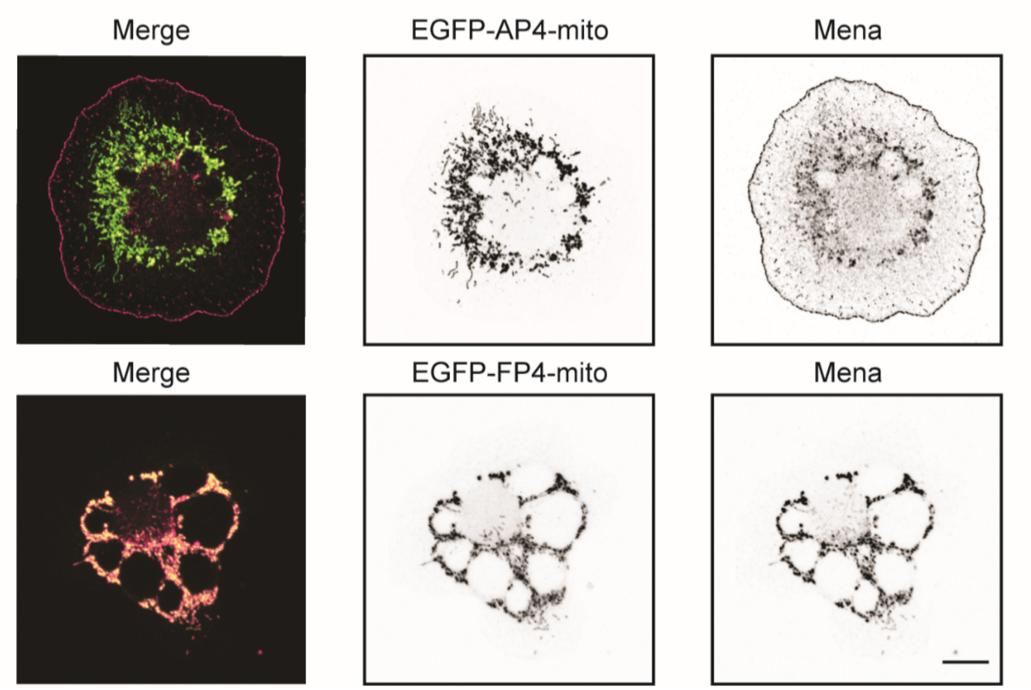


**Figure S2. Expression of EGFP-FP4-Mito construct sequesters Mena/VASP, related to Figures 1,3.** Representative images of PFN1 KO cells expressing EGFP-FP4-mito or EGFP-AP4-mito and immuno-stained for Mena. Scale bar is 10 μm.

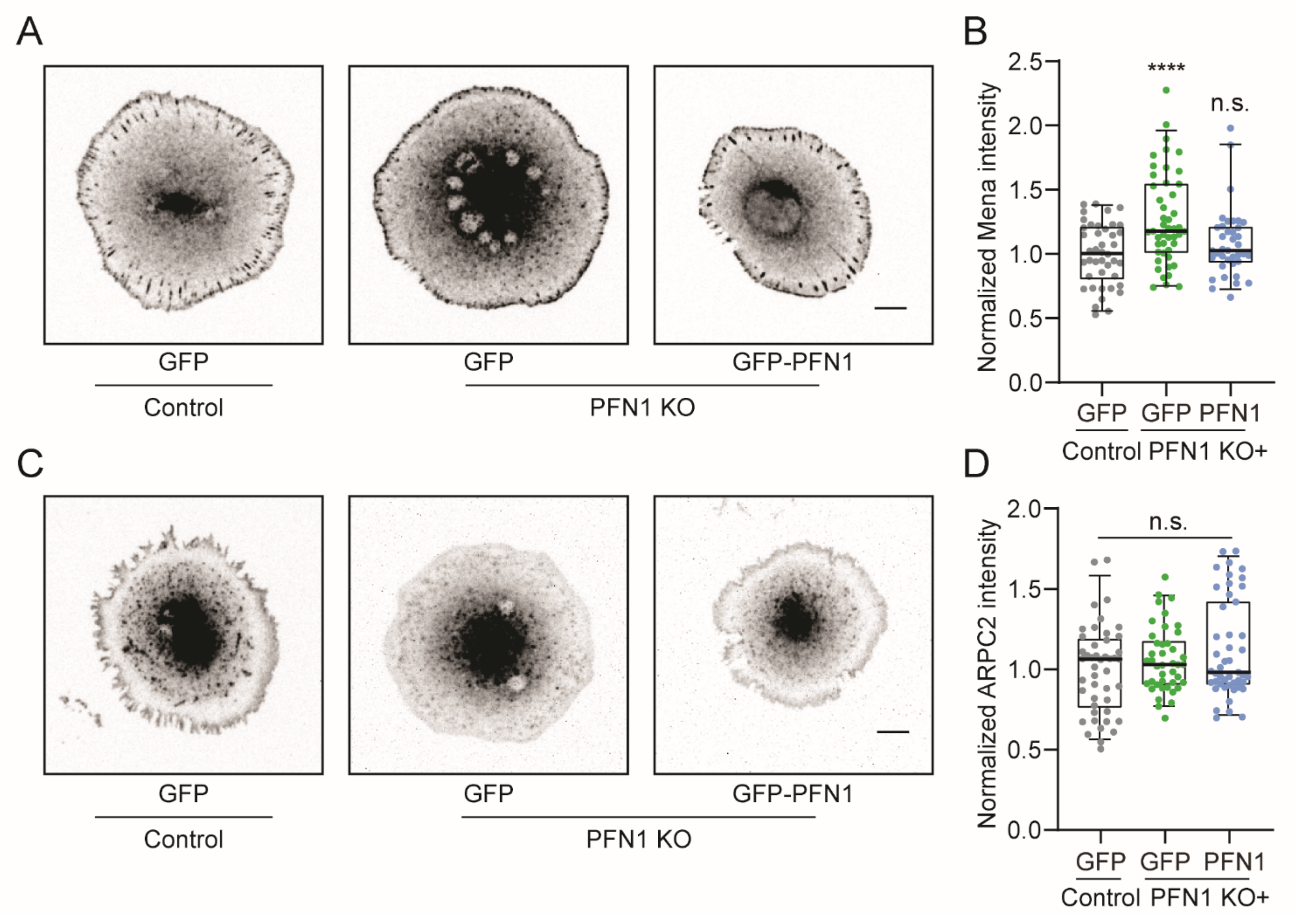


**Figure S3. Quantification of Mena and Arp2/3 expression in control and PFN1 KO cells, related to Figure 3. (A)** Images of control and PFN1 KO cells transfected with GFP or GFP-PFN1, and immunostained for Mena. **(B)** Quantification of Mena fluorescence intensity (n = 48 for control + GFP; n = 53 for PFN1 KO + GFP; n = 54 for PFN1 KO + GFP-PFN1). **(C)** Images of control and PFN1 KO cells transfected with GFP or GFP-PFN1, and immunostained for ARPC2. **(D)** Quantification of ARPC2 fluorescence intensity (n = 47 for control + GFP; n = 43 for PFN1 KO + GFP; n = 48 for PFN1 KO + GFP-PFN1). Scale bars are 10 μm.

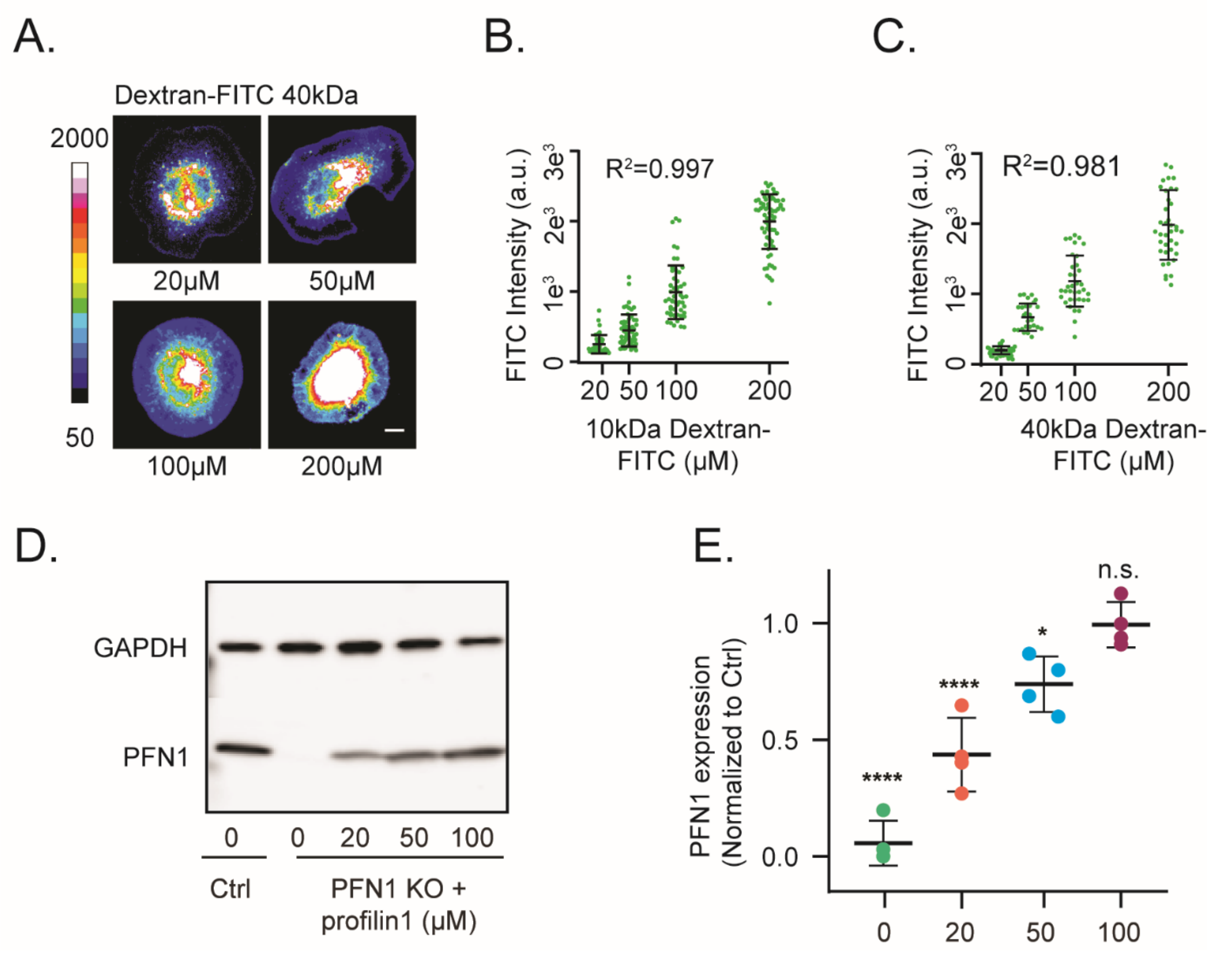


**Figure S4. Quantitative delivery of protein into cells by electroporation. (A)** Representative images of cells electroporated with either 10 or 40 kDa Dextran-FITC. The Dextran-FITC bath concentration in the electroporation chamber is indicated. The images are scaled identically and pseudocolored based on the included lookup table to convey relative fluorescent intensities. Scale bar, 10 μm. (**B,C)** Quantification of mean cellular Dextran-FITC fluorescence as a function of its bath concentration in the electroporation chamber. Individual data points are plotted along with the mean and 95% confidence intervals. The $R^2$ value of the linear fit through the mean value of each bath concentration. For 10kDa Dextran FITC **(B)** n = 42, 59, 53, 69 for 20 μM, 50 μM, 100 μM, 200 μM, respectively. For 40kDa Dextran FITC **(C)** n = 42, 27, 36, 39 for 20 μM, 50 μM, 100 μM, 200 μM, respectively. **(D)** Western blot of profilin 1 in control and PFN1 KO cells after electroporation with the designated concentration of purified profilin 1. The concentrations reflect the bath concentration of profilin 1 in the electroporation chamber. **(E)** Quantification of profilin 1 expression levels from **(D)**. Profilin 1 expression was normalized to GAPDH. Four biological replicates were used for each condition. Individual data points are plotted along with the mean and 95% confidence intervals.
